# Sexual dimorphism in gene expression and regulatory networks across human tissues

**DOI:** 10.1101/082289

**Authors:** Cho-Yi Chen, Camila Lopes-Ramos, Marieke L. Kuijjer, Joseph N. Paulson, Abhijeet R. Sonawane, Maud Fagny, John Platig, Kimberly Glass, John Quackenbush, Dawn L. DeMeo

## Abstract

Sexual dimorphism manifests in many diseases and may drive sex-specific therapeutic responses. To understand the molecular basis of sexual dimorphism, we conducted a comprehensive assessment of gene expression and regulatory network modeling in 31 tissues using 8716 human transcriptomes from GTEx. We observed sexually dimorphic patterns of gene expression involving as many as 60% of autosomal genes, depending on the tissue. Interestingly, sex hormone receptors do not exhibit sexually dimorphic expression in most tissues; however, differential network targeting by hormone receptors and other transcription factors (TFs) captures their downstream sexually dimorphic gene expression. Furthermore, differential network wiring was found extensively in several tissues, particularly in brain, in which not all regions exhibit strong differential expression. This systems-based analysis provides a new perspective on the drivers of sexual dimorphism, one in which a repertoire of TFs plays important roles in sex-specific rewiring of gene regulatory networks.

**Highlights:**

1. Sexual dimorphism manifests in both gene expression and gene regulatory networks
2. Substantial sexual dimorphism in regulatory networks was found in several tissues
3. Many differentially regulated genes are not differentially expressed
4. Sex hormone receptors do not exhibit sexually dimorphic expression in most tissues

## Introduction

Sexual dimorphism refers to the phenotypic difference between males and females of the same species (Poissant et al., 2010; Williams and Carroll, 2009). These differences present themselves in the characteristics of morphology, physiology, psychology and behavior. Physiological differences primarily exist in sex organs and reproductive systems, but also manifest in other systems, including the musculoskeletal, respiratory, and nervous systems (Blecher and Erickson, 2007; Morris et al., 2004).

Sexual dimorphism is also prevalent in human diseases. A wide range of diseases present differently in females and males, including atherosclerosis, diabetes, osteoporosis, asthma, neuropsychological disorders, and autoimmune diseases (Kaminsky et al., 2006; Morrow, 2015; Ober et al., 2008). For example, systemic lupus erythematosus is an autoimmune disease predominantly occurring in females, in a ratio of 9:1 female-to-male (Lisnevskaia et al., 2014). Differences in incidence, prevalence, severity, and response to treatment between the sexes can complicate our understanding and hinder our ability to cure and prevent diseases. Phenotypic differences between sexes may have a genetic basis (Ober et al., 2008; Poissant et al., 2010; Williams and Carroll, 2009). However, the extent to which genetics drives these effects and the mechanisms responsible are not yet fully understood (Morrow, 2015), partly because the sex-related differences in gene expression in the autosomes are usually subtle and difficult to detect. A systems-based analysis that integrates multi-omics data and collected from a large cohort of research subjects will advance our understanding of the molecular basis of sexual dimorphism.

We conducted a comprehensive expression and network analysis of 8,716 human transcriptomes collected from 28 solid tissues, whole blood, and two cell lines using the Genotype-Tissue Expression (GTEx) data set version 6.0. We compared autosomal gene expression between males and females for each tissue and found that sexual dimorphism in gene expression was seemingly ubiquitous across a wide range of human tissues. However, both the variability in expression and the number of genes involved were tissue-dependent. Gene expression of sex hormone receptors was not differentially expressed between males and females in most tissues, implying that downstream sexually dimorphic regulation is not solely explained by differences in mRNA levels of these receptors. We used a network modeling approach (PANDA+LIONESS) to infer sample-specific gene regulatory networks (Glass et al., 2013; Kuijjer et al., 2015) and compared the networks between males and females. We found significant differences in the structure of gene regulatory networks between males and females in several tissues. This suggests that differences in gene regulatory networks and their associated processes could be a characteristic of sexual dimorphism, and that changes in gene regulatory networks may explain not only differences in tissue expression, but may also help us to understand the mechanism of sexual dimorphism in diseases and other complex traits.

## Results

### Sexual dimorphism in autosomal gene expression

An overview of this study is shown in Figure 1A. RNA-Seq data from GTEx version 6.0 release were downloaded from dbGaP, preprocessed and normalized as outlined in the supplemental methods and described in more detail in (Paulson et al.). We examined only tissues with samples collected from both men and women, which included 8,716 total samples covering 31 tissues (including 28 solid organ tissues, whole blood, and two derived cell lines) from 549 research subjects (Table S1 and Table S2). Figure 1B shows the demographic information for the 549 subjects included in our analysis.

**Figure 1.**
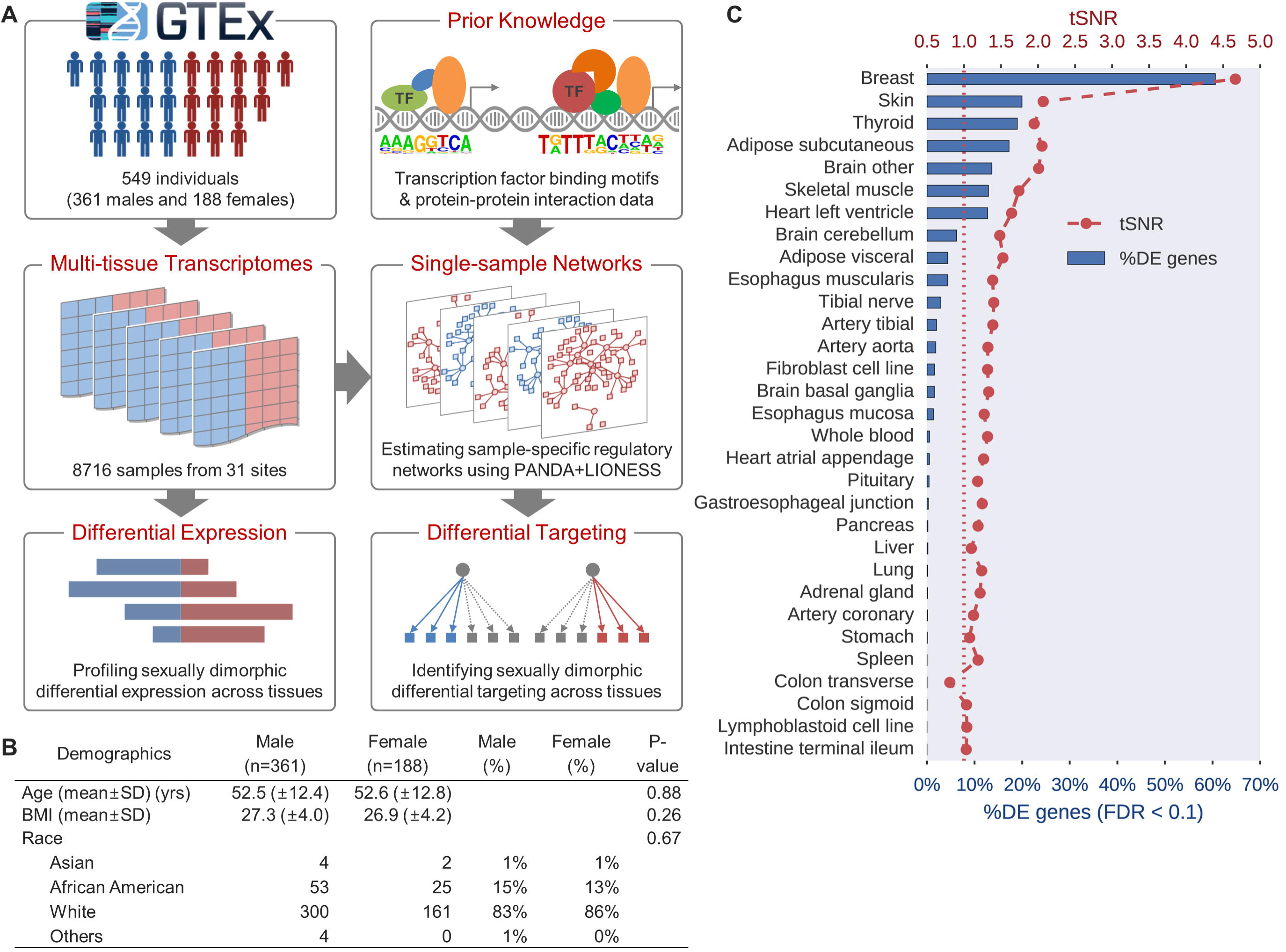
Overview of this study. **(A)** The study workflow. The GTEx v6 data set was used and annotated based on GENCODE vl9. **(B)** Demographics of the 549 study subjects. The “Others” class includes 2 American Native or Alaska Native subjects and 2 subjects of unknown race. P-values were calculated using *t*-test (for age and BMI) and Chi-square test (for race). BMI is the abbreviation of Body Mass Index (kg/m^2^). **(C)** Male-vs-female transcriptomic distance (tSNR) and percentage of differentially expressed (DE) genes in the genome (at FDR < 0.1) were used to quantify sexual dimorphism in gene expression across 31 tissues. Brain other: cerebral cortex and a set of associated structures.

We used voom (Law et al., 2014) to identify autosomal genes that were differentially expressed (DE) between males and females at a false discovery rate (FDR) less than 0.1 for each tissue (referred to as the sexually dimorphic DE genes) (Materials and Methods; File S1). We also defined a transcriptomic signal-to-noise ratio (tSNR) as a measure of the overall distance between male and female transcriptomes. For the tSNR, the signal was defined as the Euclidean distance of average gene expression profiles between groups (in this case, females and males), and the noise was defined as the overall variation among individuals (Materials and Methods). The tSNR measures the overall divergence of transcriptomes between males and females while the proportion of DE genes focuses on gene-specific sexually dimorphic expression.

An overview of sexually dimorphic gene expression is shown by tissue in Figure 1C. We found the tSNR and the proportion of DE genes are highly correlated (Pearson's *r* = 0.98) across tissues. We observed sexually dimorphic gene expression across autosomes in most tissues, with 22/31 tissues having tSNRs significantly higher than expected by chance (*P* < 0.05, permutation test) (Table S3). Breast, skin, thyroid, brain, and adipose tissues were the most sexually dimorphic (more than 10% of autosomal genes were DE), whereas the gastrointestinal tract is the least (only two genes were DE).

Tissues that are anatomically close or compositionally similar also demonstrated differing levels of sexually dimorphic expression. For example, subcutaneous adipose tissue exhibited greater sexually dimorphic gene expression than visceral adipose tissue (17% vs 4% of autosomal genes were DE, respectively). In the brain, the cerebral cortex and related structures (Other Brain) were more sexually dimorphic than cerebellum and basal ganglia (14%, 6%, and 2% of autosomal genes were DE, respectively).

We performed a pre-ranked Gene Set Enrichment Analysis (GSEA) based on the weighted *t-* statistics derived from the differential expression analysis to identify sex-biased enrichment of biological processes and associations of diseases and phenotypes in the sexually dimorphic gene signature in each tissue (Materials and Methods). We found a great diversity of functional gene sets enriched across tissues (File S2-S3). For example, energy metabolism-related genes were enriched in the adipose tissues, while the neurotransmitter secretion/transport and immune response-related gene sets were enriched in brain (Figure S1).

Additionally, disease and phenotype associations were highly enriched in a tissue-specific manner. For example, in brain, gene sets associated with neurodegenerative disorders and immune-related diseases were enriched in the brain sexually dimorphic gene signature; in heart, gene sets associated with heart block, syncope, ventricular arrhythmia, atrial fibrillation, and palpitations were enriched; in lung, we found enrichment for genes associated with chronic obstructive pulmonary disease and bronchial abnormalities; and in liver, we found ascites and hepatitis-associated gene sets. Additional findings are detailed in Files S2-S3. Overall, our results suggest that the genes with sexually dimorphic expression patterns may be involved in biological processes linked to human diseases and phenotypes, which may help explain sexual dimorphism in development and disease.

### A core set of sexually dimorphic genes shared by multiple tissues

We used Fisher's method to combine the results from the sex-specific differential expression analyses to find a common set of genes that share sexually dimorphic expression patterns across multiple tissues (Materials and Methods). We identified a total of 1,568 genes as sexually dimorphic in multiple tissues (FDR < 0.1) (Figure 2A; File S4). Many of the top significant genes, such as *FRG1B, DDX43, SPESP1, NOX5,* and *OOEP*, have been linked to sex-specific functions: *FRG1B* is a putative target gene of human androgen receptor; *SPESP1* encodes sperm equatorial segment protein involved in sperm-egg binding and fusion (Wolkowicz et al., 2008); NOX5 regulates redox-dependent processes in lymphocytes and spermatozoa (El Jamali et al., 2008); OOEP is a component of a subcortical maternal complex which was found essential during embryonic development in a mouse model (Tashiro et al., 2010). *OOEP, DDX43, SPESP1*, and *NOX5* are all highly expressed in human testis.

**Figure 2.**
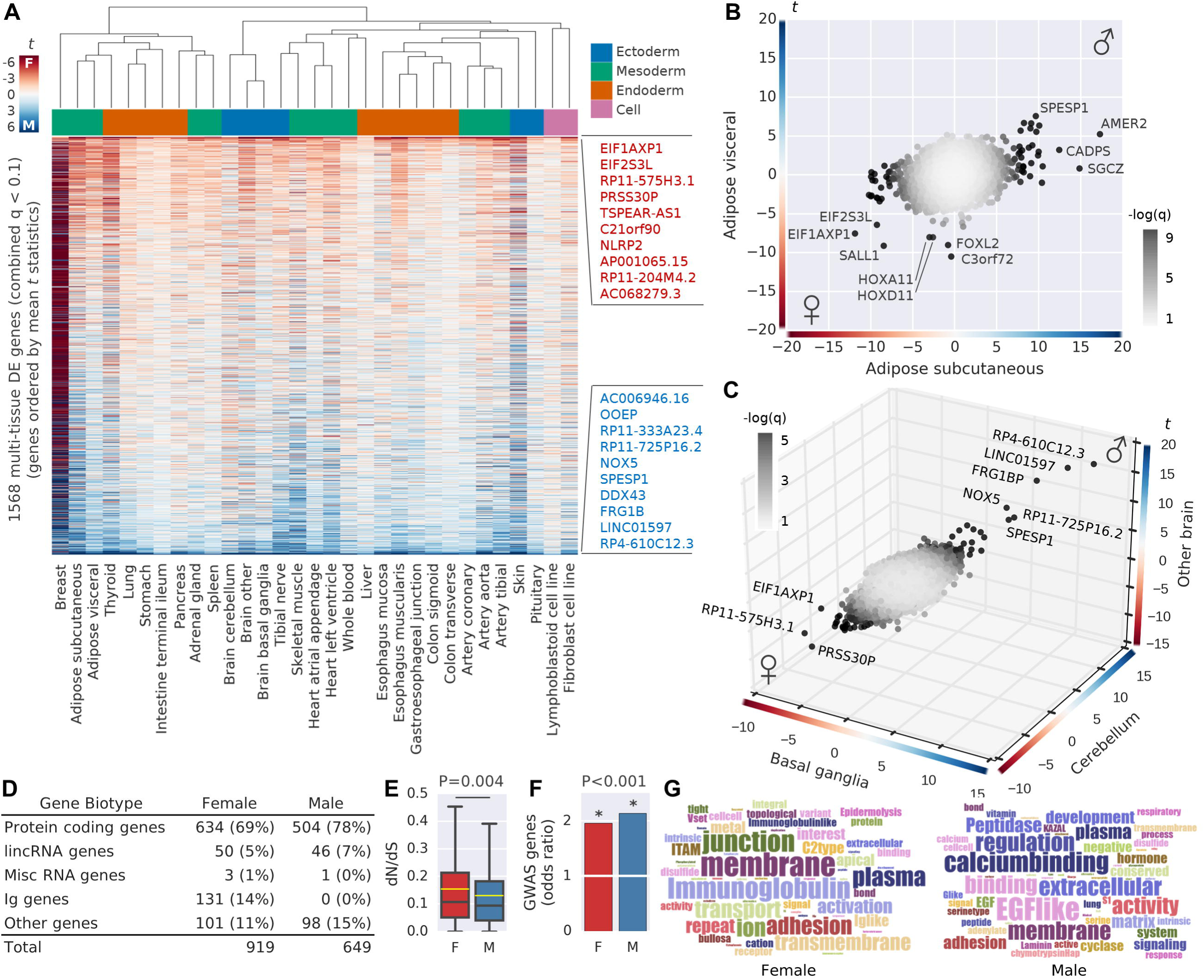
Multi-tissue sexually dimorphic DE genes. **(A)** Hierarchical clustering of the top 1568 genes with tissue-wide sexually dimorphic gene expression (combined q < 0.1 under Fisher's method). The color code of heat map depicts the *t*-statistics of sexually dimorphic differential expression. The color bar on top of the heatmap depicts germ layers of origin. Annotated on the right are the top 10 female- or male-biased DE genes. **(B-C)** Comparison of *t*-statistics of sexually dimorphic differential expression for all autosomal genes between two adipose tissue types **(B)** and between three brain clusters **(C).** *t* > 0: overexpression in males, depicted in shades of blue; *t* < 0: overexpression in females, depicted in shades of reds. Grey shades represent the significance level of data points (combined q-value). **(D)** Composition of gene biotypes of these 1568 multi-tissue sexually dimorphic DE genes. Female: genes overexpressed in females. Male: genes overexpressed in males. **(E-F)** Characteristics of the protein coding genes in the list of 1568 top multi-tissue sexually dimorphic DE genes. **(E)** Distributions of dN/dS ratio against mouse homologs. P-values were calculated using two-sample *t*-test. **(F)** Enrichment of GWAS disease-associated genes. P-values were calculated using Fisher's Exact Test (*: p < 0.001). **(G)** Word clouds of enriched Gene Ontology terms. Word size reflects term frequency.

The expression patterns of the 1,568 multi-tissue sexually dimorphic genes showed similarities between anatomically close or compositionally comparable tissue sites (Figure 2A). For example, subcutaneous adipose tissue was clustered with visceral adipose tissue; different brain regions were also clustered together. Closer inspection shows that the two adipose sites shared similar sexually dimorphic gene signature (Figure 2B); this was also observed in the brain regions (Figure 2C). These results suggest that the sub-regions of the same primary tissue often share similar sexually dimorphic gene expression patterns for a core set of genes.

We were also interested in whether there were characteristics of sexually dimorphic genes specific to each of the sexes. We categorized the 1,568 multi-tissue sexually dimorphic genes into female- (N = 919) or male-biased genes (N = 649) based on their mean *t-statistics* across all the tissues (Figure 2D). We found male-biased genes are under higher negative selective pressure (lower dN/dS ratio) than female-biased genes (*P* = 0.004, two-sample *t-*test) (Figure 2E). Both male- and female-biased genes were overrepresented as human genome-wide association (GWAS) catalog disease genes (female, *P* = 1.67 x 10^−16^; male, *P =* 7.96 x 10^−17^) (Figure 2F). Many immunoglobulin genes were female-biased (Figure 2D & 2G), consistent with previous reports (Bouman et al., 2005; Gonzalez-Quintela et al., 2008; Verthelyi, 2001). This is consistent with the enrichment of sexually dimorphic DE genes involved in immune response processes and associated with autoimmune diseases, such as rheumatoid arthritis, lupus erythematosus, ulcerative colitis, multiple sclerosis, and atopic dermatitis, that we noted previously.

As reported by others, we found long intergenic noncoding RNAs (lincRNAs) to be sexually dimorphic (Chen et al., 2012; Mele et al., 2015; Melia et al., 2015). Among the 1,568 multi-tissue sexually dimorphic genes, 50 (5%) of the female-biased genes and 46 (7%) of male-biased genes encode long intergenic noncoding RNAs (lincRNAs). Several of them were the most strongly tissue-wide sexually dimorphic genes. For example, *RP4-610C12.3* and *RP4-610C12.4* (*LINC01597)* were highly male-biased across tissues; *RP11-575H3.1* was highly female-biased (Figure 2A).

### Differential network targeting mediated by estrogen receptors

The gene-expression differences between men and women that we observed likely represent distinct, sex-specific gene regulatory processes (Williams and Carroll, 2009). One of the candidate drivers of these differences could be estrogen. The estrogen receptor (ER) is a ligand-inducible TF that can be activated by estrogen and then regulates the gene expression of a large number of target genes through binding to specific palindromic DNA sequences called estrogen response elements (EREs) (Jin et al., 2004; Welboren et al., 2007). By orchestrating the transcription of target genes, ERs mediate the physiological effects of estrogen in a diverse range of mammalian tissues and thus play an important role in growth, development, reproduction, immunology, mental health, and human disease (Au et al., 2016; Hara et al., 2015).

Given the important role of estrogen and ER in sexual development and reproductive function, we might expect the gene expression of ER to be sexually dimorphic. However, both estrogen receptor genes, estrogen receptor alpha (*ESR1*) and beta (*ESR2*), were not differentially expressed between males and females in most of the tissues we analyzed, including highly sexually dimorphic tissues such as breast, brain, and muscle (Figure S2). This suggests that estrogen may exert its sexually dimorphic effects by altering gene regulatory networks without differential expression of its receptors.

We used PANDA+LIONESS (Glass et al., 2013; Kuijjer et al., 2015) to infer gene regulatory network models for each sample in each tissue and compared those network models between men and women (Figure 1A; Materials and Methods). Briefly, we applied PANDA to create a consensus network model that comprises 652 TFs and 27,175 target genes by integrating multiple sources of data including gene expression profiles (GTEx), TF binding motifs (CIS-BP) (Weirauch et al., 2014), and protein-protein interactions (StringDB) (Szklarczyk et al., 2015). We then used LIONESS to infer individual networks for each sample in the population via linear interpolation. PANDA+LIONESS produced 8,716 inferred gene regulatory network models, one for each RNA-Seq transcriptome from the 549 GTEx research subjects. These network models represent the strength of the estimated regulatory interaction between TFs and their target genes as weighted bipartite directed graphs, in which nodes represent TFs or their target genes and edges represent inferred regulatory relationships between nodes.

Analogous to how we identified sexually dimorphic DE genes, we used limma to compare the weight of each edge between males and females in each tissue and identified “differentially targeting” edges, which represent a set of TF-target relationships that exhibit differential regulatory potential between the sexes (File S5; Materials and Methods). This allowed us to identify regulatory network “wiring patterns” that differed between males and females in each tissue and to identify key regulators that may contribute to the shaping of the sexually dimorphic gene expression landscape in each tissue.

We started by investigating estrogen receptors in the breast tissue. Although the estrogen receptors *ESR1* and *ESR2* were not differentially expressed between males and females, we found that the regulatory targeting by these receptors was different between the sexes. We highlight two of the most sexually dimorphic DE genes that are targets of ESR1 in our network models, *IGLV4-69* and *MYOT,* as examples. *IGLV4-69* is an immunoglobulin gene that had increased expression in females yet increased targeting by ESR1 in males (Figure 3A). In contrast, *MYOT*, a gene that encodes a cytoskeletal protein (myotilin) that stabilizes thin filaments during muscle contraction, showed over-expression in males but greater ESR1-targeting in females (Figure 3B). In both cases, edge weights connecting ESR1 to its target genes linearly reflected the expression of these target genes (Figure 3C-D, upper). However, *ESR1* itself was not differentially expressed between males and females, and its expression poorly correlated with its target gene expression (Figure 3C-D, bottom). This implies that the edge weights in the network models capture the regulatory activities of ESR1 while the gene expression of *ESR1* alone does not.

**Figure 3.**
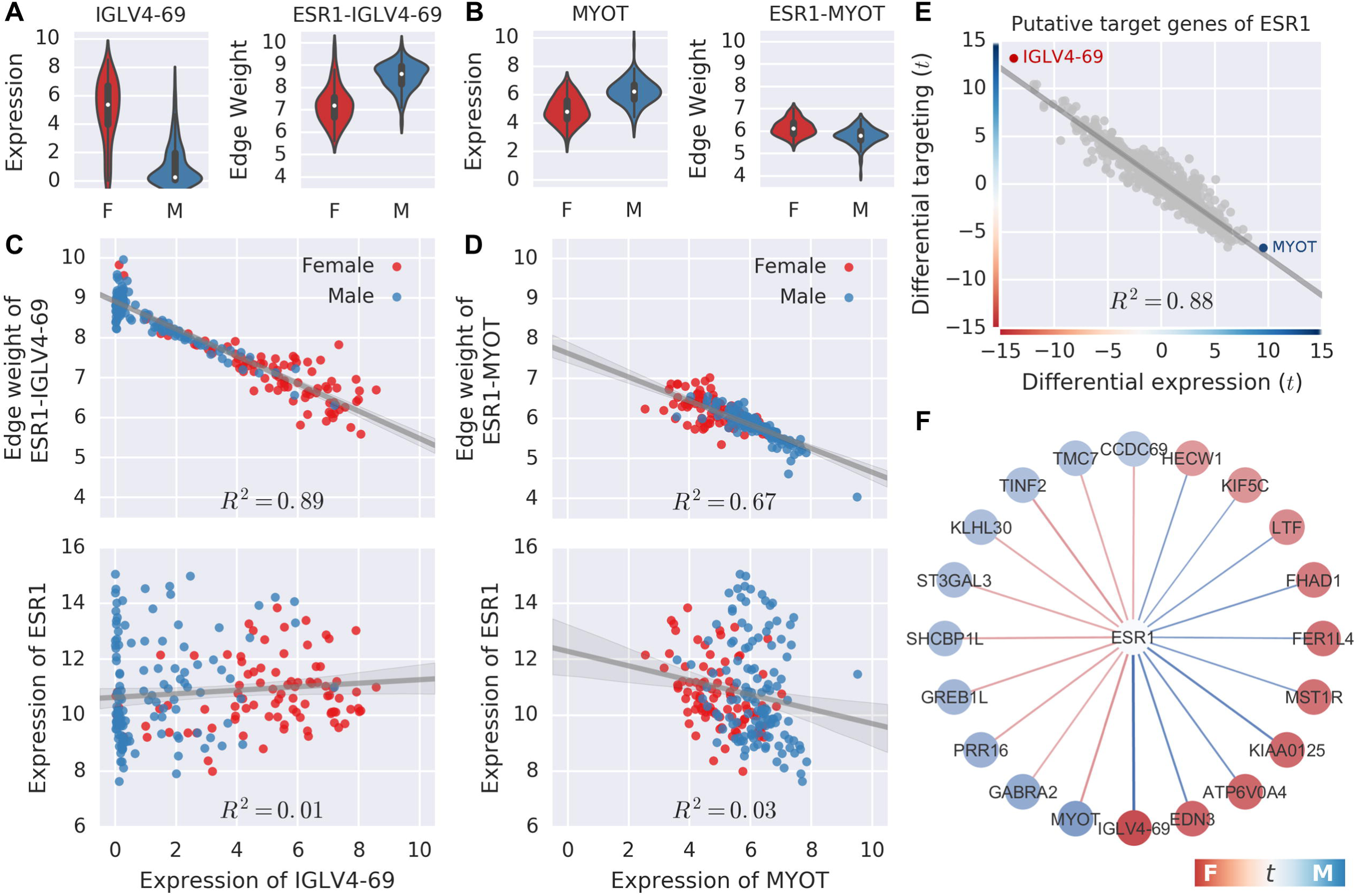
Network inference suggests regulatory potential of ESR1 in driving the sexually dimorphic gene expression of its target genes. **(A-B)** Comparison of gene expression levels and targeting levels by ESR1 between females and males in breast for *IGLV4-69* **(A)** and *MYOT* **(B). (C-D)** Expression of *IGLV4-69* **(C)** and *MYOT* **(D)** are highly correlated with targeting levels by ESR1 (upper) but not with *ESR1* expression (bottom). Each data point represents a single sample. **(E)** Regression between differential expression levels and differential targeting levels by ESR1 on its putative targets. **(C-E)** R^2^: adjusted coefficient of determination. **(F)** Network visualization of the top 20 genes significantly differentially targeted by ESR1. Edge color represents the direction and strength of differential targeting. Node color represents the direction and strength of differential expression.

By considering the putative set of ESR1 target genes in the breast tissue, we found that the sexually dimorphic differential targeting by ESR1 explains much of the variance (adjusted R^2^ = 0.88) in the differential expression of its targets (Figure 3E), suggesting that the regulatory influence of ESR1 may drive the sexually dimorphic gene expression of its target genes. We note that ESR2 shows a similar regulatory pattern in the regulatory networks in the breast tissue (adjusted R^2^ = 0.81) (Figure S3).

Figure 3F shows the 20 genes most differentially targeted by ESR1 in the breast network models. Several of these genes are implicated in breast function and disease. For example, *MST1R*, encodes a macrophage stimulating receptor associated breast cancer progression (Privette Vinnedge et al., 2015). *ATP6V0A4* mediates invasion of breast cancer cell lines in the transwell Matrigel assay (Hinton et al., 2009). *EDN3* has been suggested as a tumor suppressor gene in the human mammary gland (Wiesmann et al., 2009) and its expression was shown to be regulated by estrogen and progesterone in the rhesus macaque (Keator et al., 2011). *LTF* (Lactotransferrin) is an estrogen-inducible gene (Das et al., 1998; Ghosh et al., 1999; Moggs et al., 2004; Stuckey et al., 2006); its protein product exists in high concentration in human milk (Hakansson et al., 1995).

### A repertoire of TFs modulate sexually dimorphic gene expression

To investigate whether the targeting of genes by ESRs is equally important in tissues other than breast, and to investigate if other TFs also drive sexually dimorphic gene expression, we extended our analysis to include all 652 TFs and analyzed them in each tissue. We found that ESR1 influences gene expression differently across tissues. In subcutaneous adipose and thyroid, for example, the differential targeting of ESR1 largely explains the sexually dimorphic gene expression of its target genes (adjusted R^2^ = 0.78 and 0.59, respectively) (Figure 4A-B), whereas in colon the relationship was relatively poor (adjusted R^2^ = 0.18) (Figure 4C).

**Figure 4.**
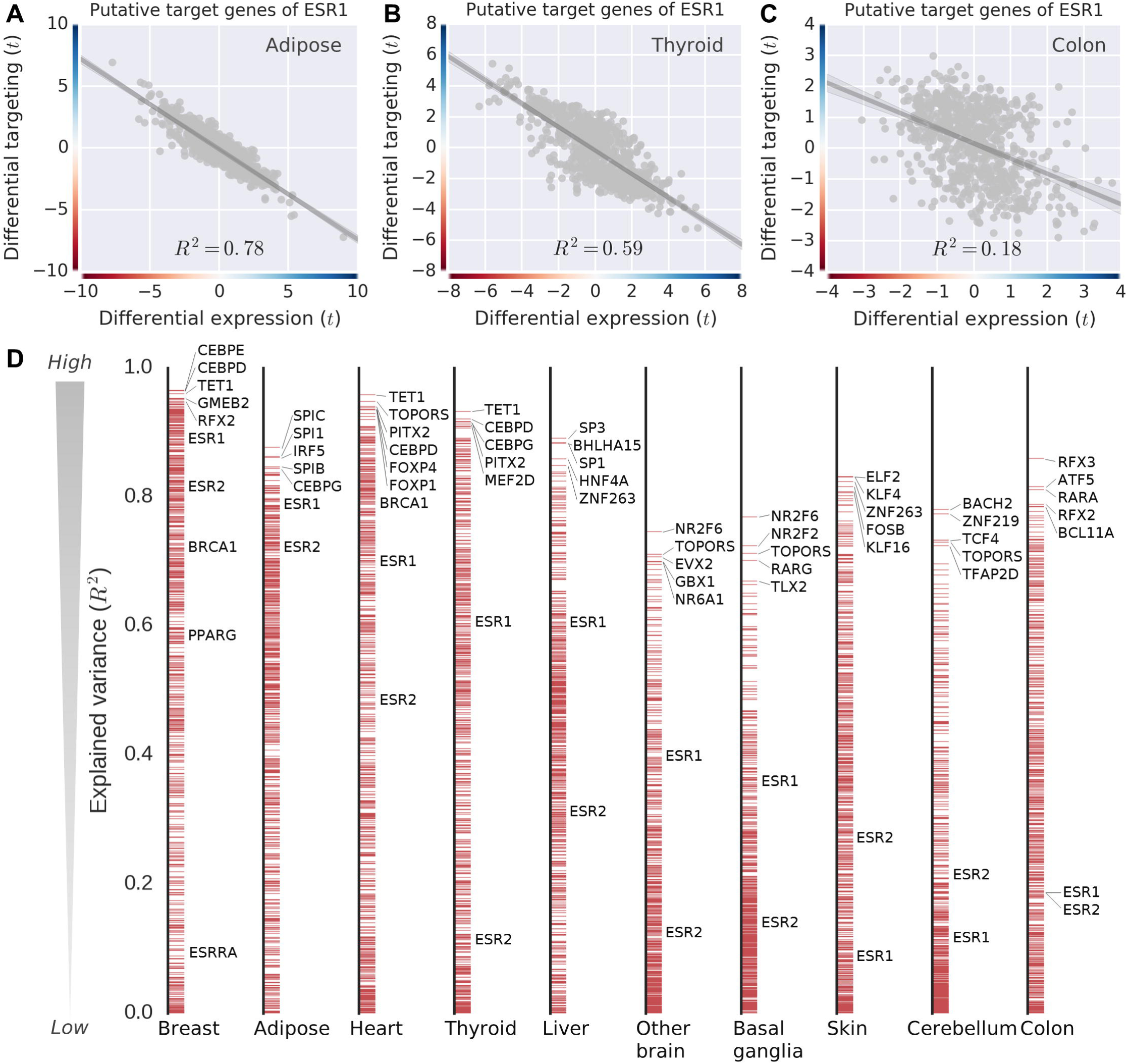
Key transcription factors that shape the sexually dimorphic gene expression landscape. **(A-C)** Inferring differential targeting of ESR1 to its putative target genes versus differential expression levels in three representative tissues. **(D)** Adjusted coefficient of determination (R^2^) was used to quantify and rank the regulatory potential of 652 TFs in driving the sexually dimorphic gene expression of their target genes. Positions of representative TFs are highlighted approximately in the plot. See Table 1 for TFs' documented roles in hormone-responsive pathways and sex-related biological processes and diseases. Adipose: subcutaneous adipose tissue. Fleart: left ventricle. Brain other: cerebral cortex and a set of associated structures. Colon: sigmoid colon.

By ranking all 652 TFs in our network models based on how well their differential targeting explains the variance in the sexually dimorphic differential expression patterns of their target (adjusted R^2^), we found that the highest-ranking TFs in each tissue are not always hormone receptors (Figure 4D; File S6). However, the top TFs often have documented roles in hormone-responsive pathways and sex-related biological processes and diseases (Table 1). These results suggest that a wide variety of TFs mediate the sexually dimorphic regulatory programs active in different tissues.

**Table 1.** TFs that may drive downstream sexually dimorphic gene expression across tissues

| TF | Gene name | Documented roles | References |
| --- | --- | --- | --- |
| BRCA1 | Breast Cancer 1 | Responsive to estrogen; regulating acetylation and ubiquitination of ESR1; implicated in breast cancer | (Chrzan and Bradford, 2007; Crowe and Lee, 2006; Ma et al., 2010; Wang et al., 2005) |
| CEBPD/<br>CEBPE | CCAAT/enhancer binding protein, delta/epsilon | Responsive to estradiol | (Buterin et al., 2006; Tee et al., 2004) |
| FOXP1/F<br>OXP4 | Forkhead Box P1/P4 | Co-expressed with estrogen receptors; estrogen-inducible; implicated in breast and ovarian cancers; important in heart development | (Fox et al., 2004; Ijichi et al., 2013; Ikeda et al., 2015; Shigekawa et al., 2011) |
| GMEB2 | Glucocorticoid Modulatory Element Binding Protein 2 | Responsive to ethinyl estradiol | (Fong et al., 2007) |
| KLF4 | Kruppel-Like Factor 4 | Yamanaka factor (iPS factor) required for skin development; suppressing transcriptional activity of ESR1 | (Akaogi et al., 2009; Rowland and Peeper, 2006; Takahashi et al., 2007) |
| NR2F6 | Nuclear Receptor Subfamily 2 Group F Member 6 | Involved in modulation of hormonal responses and development of brain circadian clock; suppressing transcriptional activity of ESR1 induced by estrogens | (Warnecke et al., 2005; Zhu et al., 2000) |
| SP1/SP3 | Sp1/Sp3 Transcription Factor | Involved in liver development; engaged in estrogen signaling pathway via PPI with ER-ligand complexes | (Kazi et al., 2005; Moggs and Orphanides, 2001; Mukherjee et al., 2005) |
| TET1 | Tet Methylcytosine Dioxygenase 1 | Involved in DNA methylation processes; implicated in lateral myocardial infarction and coronary aneurysm | (Li et al., 2013; Rovai et al., 2016; Takai et al., 2014) |
| TOPORS | TOP1 Binding Arginine/Serine-Rich Protein | Binding to Parkinson's disease-associated protein (DJ-1) and protecting against neuronal apoptosis by regulating p53 transcriptional activity | (Shinbo et al., 2005; Xu et al., 2005) |

### Brain exhibits extensive differential network wiring

Sex differences in gene regulatory networks involve both TFs and target genes. To identify differentially targeted genes between males and females, we calculated the in-degree for each gene as the sum of its network edge weights and compared in-degrees between males and females. As shown in Figure 5A, we found a large proportion of genes were both DE and differentially targeted (DT) in several tissues including breast, thyroid, muscle, and liver.

**Figure 5.**
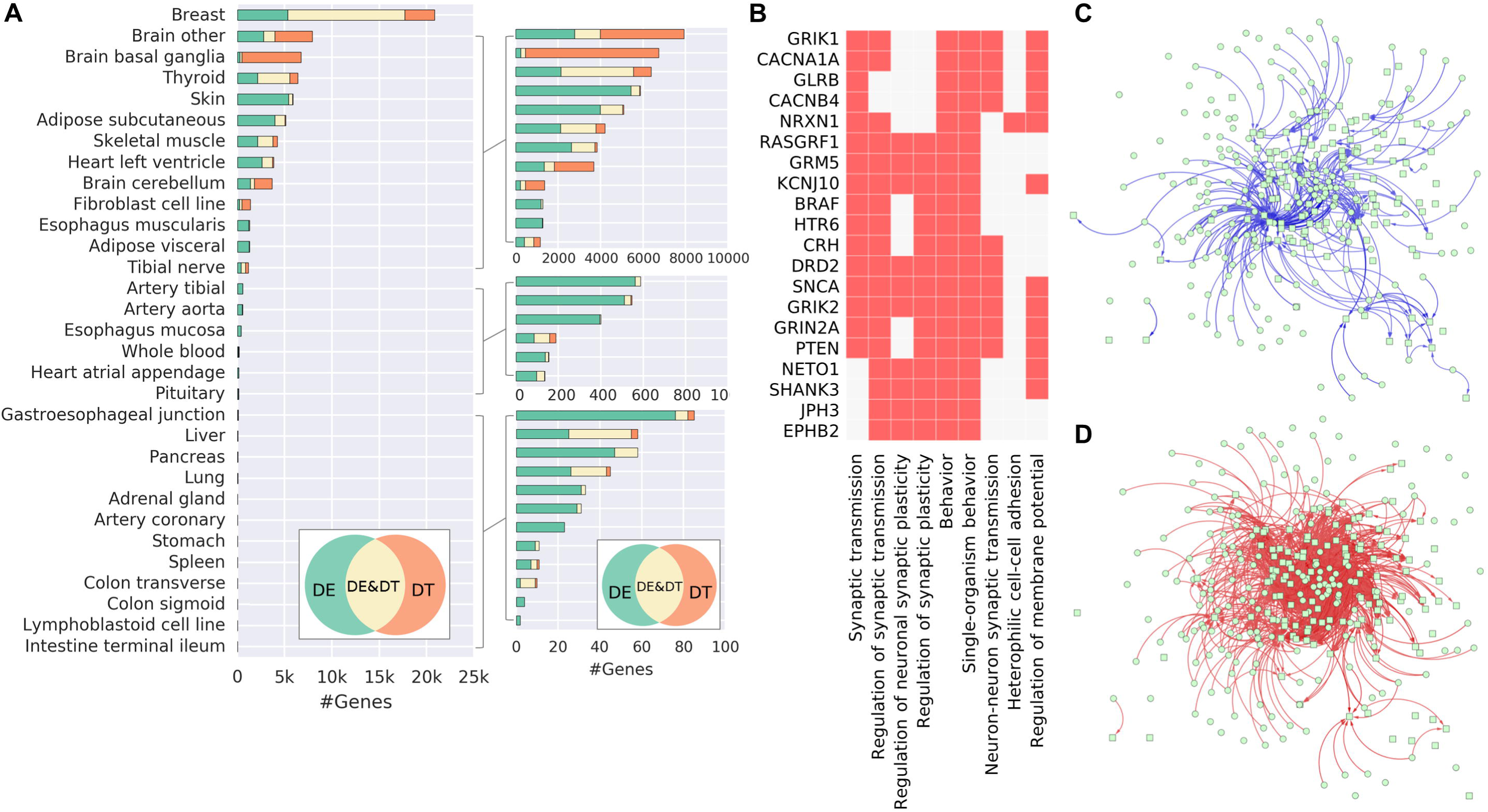
Brain networks are differentially wired between males and females. **(A)** Number of genes that are differentially expressed (DE), differentially targeted (DT), and both differentially expressed and targeted (DE&DT) across tissues. **(B)** In basal ganglia, brain functions are enriched in DT genes. X-axis: top nine enriched GO terms; y-axis: genes with shared enriched GO terms. Cells in the matrix indicate if a gene is associated with a term. **(C-D)** Visualization of the synaptic transmission subnetwork in a force-directed layout shows differential wiring in the network between males **(C)** and females **(D)** in basal ganglia.

Importantly, we also found that a surprisingly large number of genes in brain, especially in basal ganglia, are differentially targeted but not differentially expressed (6,479 DT genes at FDR < 0.1). These DT-only genes in basal ganglia are associated with brain functions, with enrichment in synaptic and behavior-related biological processes (Figure 5B). For instance, *CACNA1A*, which encodes a voltage-dependent calcium channel, has been associated with multiple diseases such as episodic ataxia (Tomlinson et al., 2016), epileptic encephalopathy (Damaj et al., 2015), and autism (Damaj et al., 2015; Li et al., 2015). As an example, we found that there are 185 TFs differentially targeting 128 genes associated with synaptic transmission process in basal ganglia between males and females (Figure 5C-D). These results reveal an additional layer of complexity of sexual dimorphism, in which differential network wiring also characterizes sex-specific gene regulatory networks.

## Discussion

Sexual dimorphism manifests in human development, physiology, and in the incidence and progression of diseases. Sexual dimorphism influences how the genetic program drives the phenotypes we observe and the ways in which those phenotypes differ between the sexes. A natural hypothesis is that the phenotypic differences between men and women in different tissues should be reflected in sex-specific gene expression in these tissues. However, a global picture of sexual dimorphism in gene expression is difficult to depict, partly because the sex-related differences in the autosomal gene expression are usually subtle and varied across tissues.

One strategy to tackle this problem is to include more samples in the analysis to increase statistical power. Our analysis was based on the GTEx version 6.0 cohort, which includes 8,716 samples from 549 individuals; we excluded the sex chromosome genes so that they would not skew estimates of sexually dimorphic expression. This large sample size enabled us to detect subtle differences in autosomal gene expression at a higher resolution than in previous studies. We discovered many tissue-specific sexually dimorphic differentially expressed genes on the autosomes, many of which have important known biological and disease-related functions.

Mele and colleagues (Mele et al., 2015) have also reported sex-biased gene expression across human tissues based on a relatively small GTEx pilot data set which included only 1,641 postmortem tissue samples from 175 individuals. They included both autosomal and sex chromosome genes and found that most sexually dimorphic DE genes were on sex chromosomes. The only exception was breast, in which 715 differentially expressed autosomal genes were found. Then they used WGCNA to identify modules of correlated gene expression and showed that those co-expression modules were enriched with GO terms that included spermatid, ectoderm, and epidermis development. However, WGCNA is a correlation-based method that captures general patterns of co-expression but does not distinguish between transcription factors and their targets, and so may fail to extract sex-specific regulatory processes.

The emergence of sex-specific traits likely involves modifications in sexually dimorphic gene expression patterns, which may be driven by sex-dependent gene regulation (Blekhman et al., 2010; Ober et al., 2008; Williams and Carroll, 2009). Ligand-dependent transcriptional regulation by steroid sex hormones and their receptors was suggested to play a role in sexual development and reproductive function by mediating downstream target gene expression with other TFs (Bouman et al., 2005; Ikeda et al., 2015; Verthelyi, 2001). To our surprise, we did not observe significant differences in gene expression of steroid hormone receptors between men and women in most of the analyzed tissues.

A hypothesis is that sexual dimorphism is a manifestation of different regulatory programs in male and female tissues. We used PANDA and LIONESS, a systems-based method that uses a prior based on TF binding motifs and integrates multi-omics data to model regulatory processes in both populations and individuals. We discovered that male and female networks have significant differences in their regulatory structure, even when the targets of specific TFs are not differentially expressed. This is a subtle but important point as tissue-specific patterns of gene expression may be maintained differentially in males and females. This means that sexual dimorphism may result from perturbations that affect the regulatory processes in one sex rather than the other, altering the expression of the downstream genes.

As an example, we presented an analysis of sexual dimorphism in the human brain. We found that the structure of gene regulatory networks differed between males and females. Differentially targeted genes in brain were enriched in biological functions and pathways relevant to brain diseases, many of which are known to exhibit differences in men and women. Indeed, sex differences have been found in brain structure, cognitive functions, behaviors, and in several brain diseases and mental disorders, including Parkinson's and Alzheimer's disease, drug abuse, anxiety, and depression (Frick and Gresack, 2003; Gillies and McArthur, 2010; McCarthy, 2008; Pol et al., 2006; Rizk-Jackson et al., 2006; Tsai et al., 2009).

Our systems-based approach to gene regulatory network modeling provides a new perspective on sexual dimorphism, one in which steroid hormones and non-steroid transcription factors affect the regulatory program by altering the structure of sex-specific gene regulatory networks. These sex-specific processes often regulate genes associated with development and disease, and may help to explain sexual dimorphism we observe. These findings across 31 tissues reiterate a phenomenon that we had observed in the study of male and female differences in chronic obstructive pulmonary disease (Glass et al., 2014).

Sexual dimorphism has been recognized as one of the most significantly understudied aspects of human disease research and the 2015 guidelines from the National Institutes of Health (NOT-OD-15-102) have identified consideration of sex as biological variable should be factored into research designs, analyses, and reporting in vertebrate animal and human studies (Health, 2015). The GTEx data provide an unprecedented opportunity to explore how sex influences gene expression. We see great diversity in the degree of sexually dimorphic gene expression, with some tissues appearing to be very different while others are almost indistinguishable. But by using PANDA and LIONESS to model regulatory processes, we find higher order differences in regulatory processes that distinguish men and women. Our analysis of 31 tissues provides an important baseline for any disease-based studies and underscores the importance of looking beyond expression to consider sexually dimorphic patterns of transcriptional regulation when studying human development and disease.

## Experimental procedures

More details are available in Supplemental Experimental Procedures.

### GTEx data set

The Genotype-Tissue Expression (GTEx) version 6.0 RNA-Seq data set (phs000424.v6.p1, 2015-10-05 released) was downloaded from dbGaP (approved protocol #9112). Using YARN in Bioconductor [bioconductor.org/packages/yarn], we performed quality control, gene filtering, and normalization preprocessing (Paulson et al.). We identified and removed GTEX-11ILO due to potential sex misannotation. We grouped related body regions using gene expression similarity. For example, skin samples from the lower leg (sun exposed) and from the suprapubic region (sun unexposed) were grouped as “skin.” We removed sex-specific tissues (prostate, testis, uterus, vagina, and ovary) and those with fewer than 30 samples of one sex (kidney cortex and minor salivary gland). The final data set contains 8716 samples from 31 tissues (which included 28 solid organ tissues, whole blood, and two derived cell lines) from 549 research subjects (188 females and 361 males) (Table S1 and Table S2). We removed sex-chromosome and mitochondrial genes (retaining 29,242 genes).

### Differential expression analysis

Differential expression (DE) analysis was performed using voom (Law et al., 2014) to transform RNA-Seq read counts to log counts per million (log-cpm) with associated precision weights, followed by linear modeling and empirical Bayes procedure using limma. In each tissue, we adopted the following linear regression model to detect sexually dimorphic gene expression:

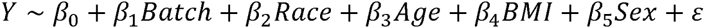

where *Y* is the gene expression; *Batch* denotes the type of nucleic acid isolation; TrueSeq RNA library preparation kit v1 was used in all GTEx RNA-Seq experiments; *Race* denotes the race of the subject; *Age* denotes the age of the subject, *BMI* denotes the body mass index of the subject, and *Sex* denotes the reported sex of the subject. The linear model is fitted to the expression data for each gene using a least-squares fitting method. The p-values for the estimated coefficient of Sex were adjusted for multiple testing using the Benjamini-Hochberg method. A false discovery rate of 0.1 was used as the significance threshold to report sexually dimorphic DE genes in each tissue. To identify differential expression across multiple tissues, we used Fisher's method (Fisher, 1970) to combine the adjusted p-values from the results of differential expression analysis in each of the GTEx tissues.

### Transcriptomic signal-to-noise ratio

Transcriptomic signal-to-noise ratio (tSNR) between expression profiles of females (*X*) and males (*Y*) was calculated as:

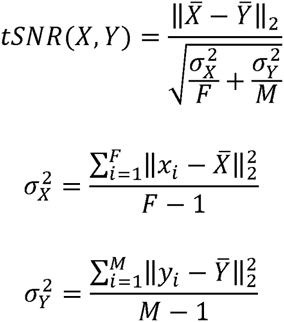

where *F* and *M* represent the sample sizes of females and males, respectively. To derive an empirical p-value for each observed tSNR in each tissue, we used a permutation test that shuffled the sex labels to samples while maintaining the total numbers of females and males unchanged for 1,000 iterations. If there was zero occurrence from the 1,000 times permutation more extreme than the observed tSNR, the empirical p-value would be set to 0.5/1,000.

### Functional enrichment analysis

Gene Set Enrichment Analysis (GSEA) was performed based on the pre-ranked list of t-statistics derived from the differential expression analysis (Subramanian et al., 2005). Figure S1 was visualized by using Enrichment map (Isserlin et al., 2014) in Cytoscape (Smoot et al., 2011). If not stated explicitly otherwise, default parameters and a default background set were used when performing the analyses. Gene annotations were obtained from the Gene Ontology Consortium (Gene Ontology, 2015), FunDO (Du et al., 2009), and Human Phenotype Ontology (HPO). Annotation files were compiled into the Gene Matrix Transposed (GMT) file format and were loaded to GSEA program (v2-2.2.2).

### Network inference using PANDA+LIONESS

Single-sample networks were reconstructed using PANDA+LIONESS (Glass et al., 2013). We performed a motif scan to generate a gene regulatory prior. We downloaded position weight matrices (PWM) for *Homo sapiens* motifs from the Catalog of Inferred Sequence Binding Preferences (CIS-BP) (Weirauch et al., 2014). We mapped the PWMs to promoter regions of Ensembl genes (GRCh37.p13) using FIMO (Grant et al., 2011). Motif mappings were parsed to only retain those below p-value cutoff of 10^−5^ and those that were within a promoter range of [-750, +250] around the transcription start site (TSS). This resulted in a regulatory prior of 652 TFs targeting 27,249 target ENSG identifiers. We generated a protein-protein interaction (PPI) prior based on the *Homo sapiens* PPI data set from StringDB version 10. We then parsed our gene expression data to match the genes in the motif prior (27,175 genes) and ran PANDA followed by LIONESS.

## Author contributions

CYC, CLR, MK, JNP, AS, MF, JP, KG, JQ, and DLD conceived the study; CYC, CLR, MK, and JNP analyzed the data; CYC, CLR, KG, JQ, and DLD interpreted the results. All authors contributed to the reviewing and editing of the manuscript. All authors read and approved the final manuscript.

## Acknowledgments

This work was supported by grants from the US National institutes of Health, including grants from the National Heart, Lung, and Blood Institute (5P01HL105339, 5R01HL111759, 5P01HL114501, K25HL133599), the National Cancer Institute (5P50CA127003, 1R35CA197449, 1U01CA190234, 5P30CA006516), and the National Institute of Allergy and Infectious Disease (5R01AI099204). Additional funding was provided through a grant from the NVIDIA foundation. This work was conducted under dbGaP approved protocol #9112 (accession phs000424.v6.p1).

## Supplemental figure legends

**Figure S1.**
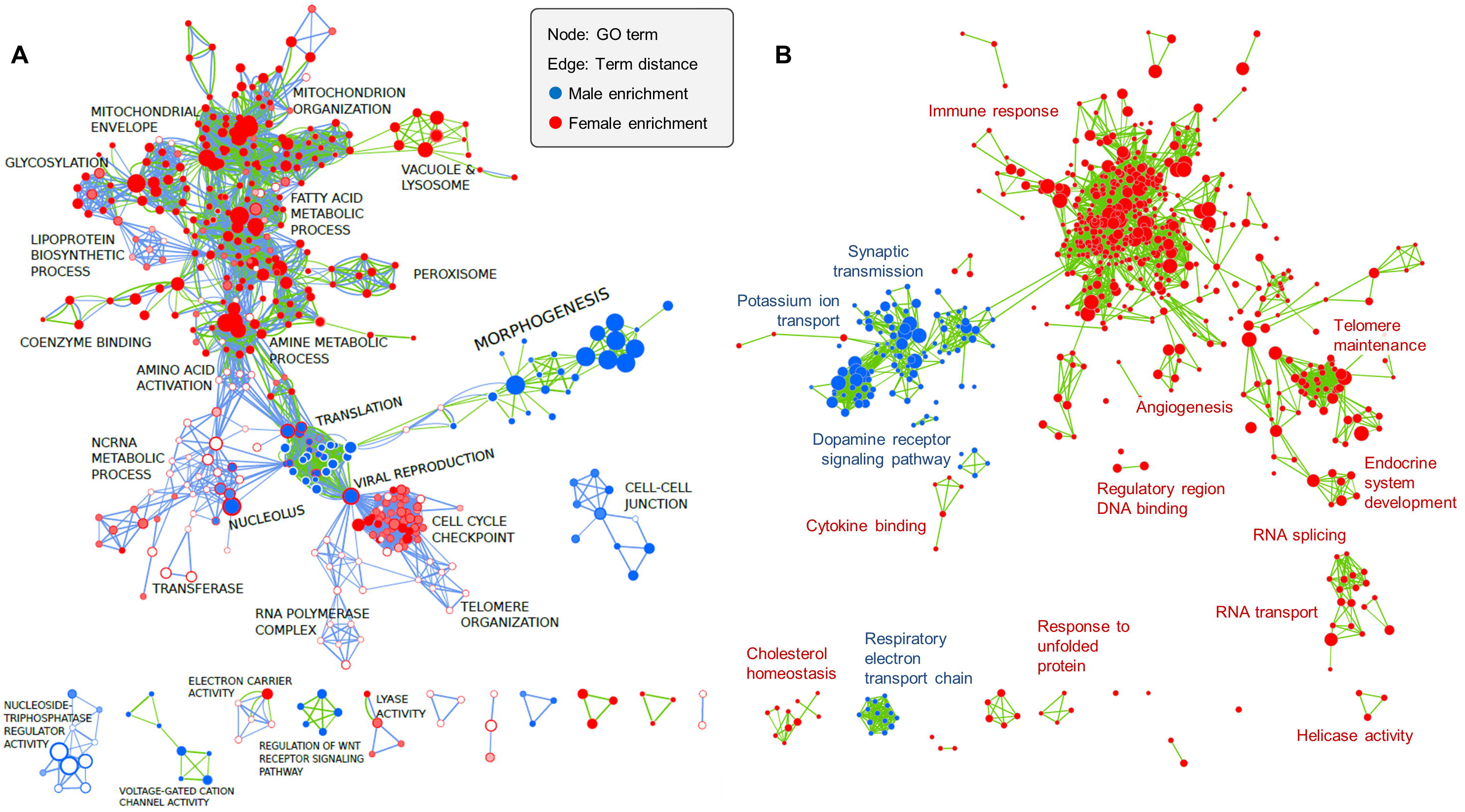
GSEA Enrichment maps for brain and adipose tissues, related to Figure 1. Maps of female-vs-male functional enrichments using GSEA based on Gene Ontology (GO) vocabulary for **(A)** adipose and **(B)** brain tissues. Nodes are GO terms (gene sets). Edges represent term distance (mutual overlap between two connected gene sets). Node size indicates gene set size. In **(A)**, filled and unfilled circles represent enrichment in subcutaneous and visceral adipose tissues, respectively. Edge colors indicate the source of enrichment: green corresponds to subcutaneous adipose tissue and blue corresponds to visceral adipose tissue.

**Figure S2.**
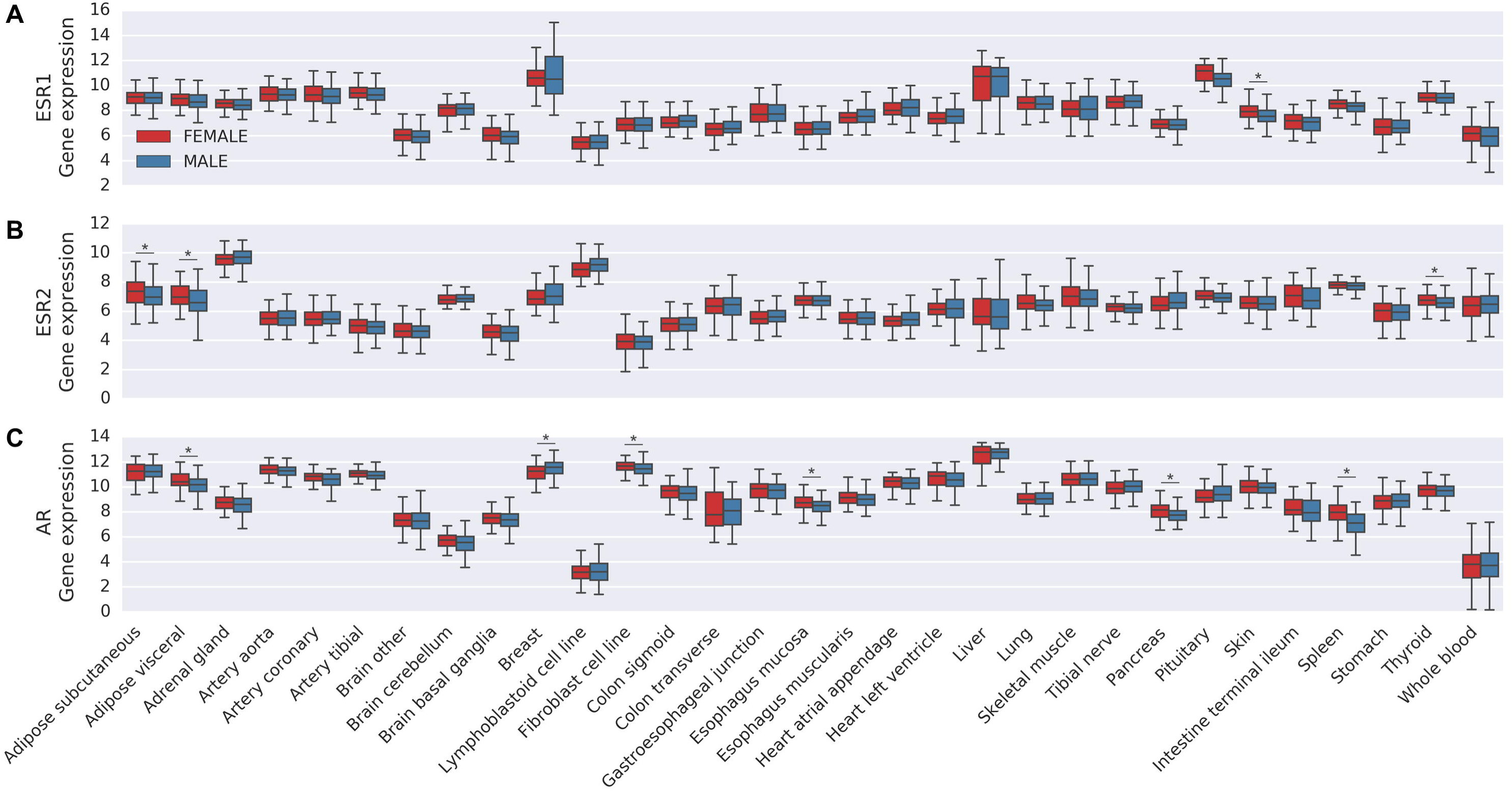
Gene expression of sex hormone receptors is not ubiquitously sexually dimorphic, related to Figure 2. **(A-C)** Distribution of mRNA expression (log-cpm, Y-axis) of **(A)** estrogen receptor alpha (ESR1), **(B)** estrogen receptor beta (ESR2), and **(C)** androgen receptor (AR) in females (red) and males (blue) across 31 sites (X-axis).

**Figure S3.**
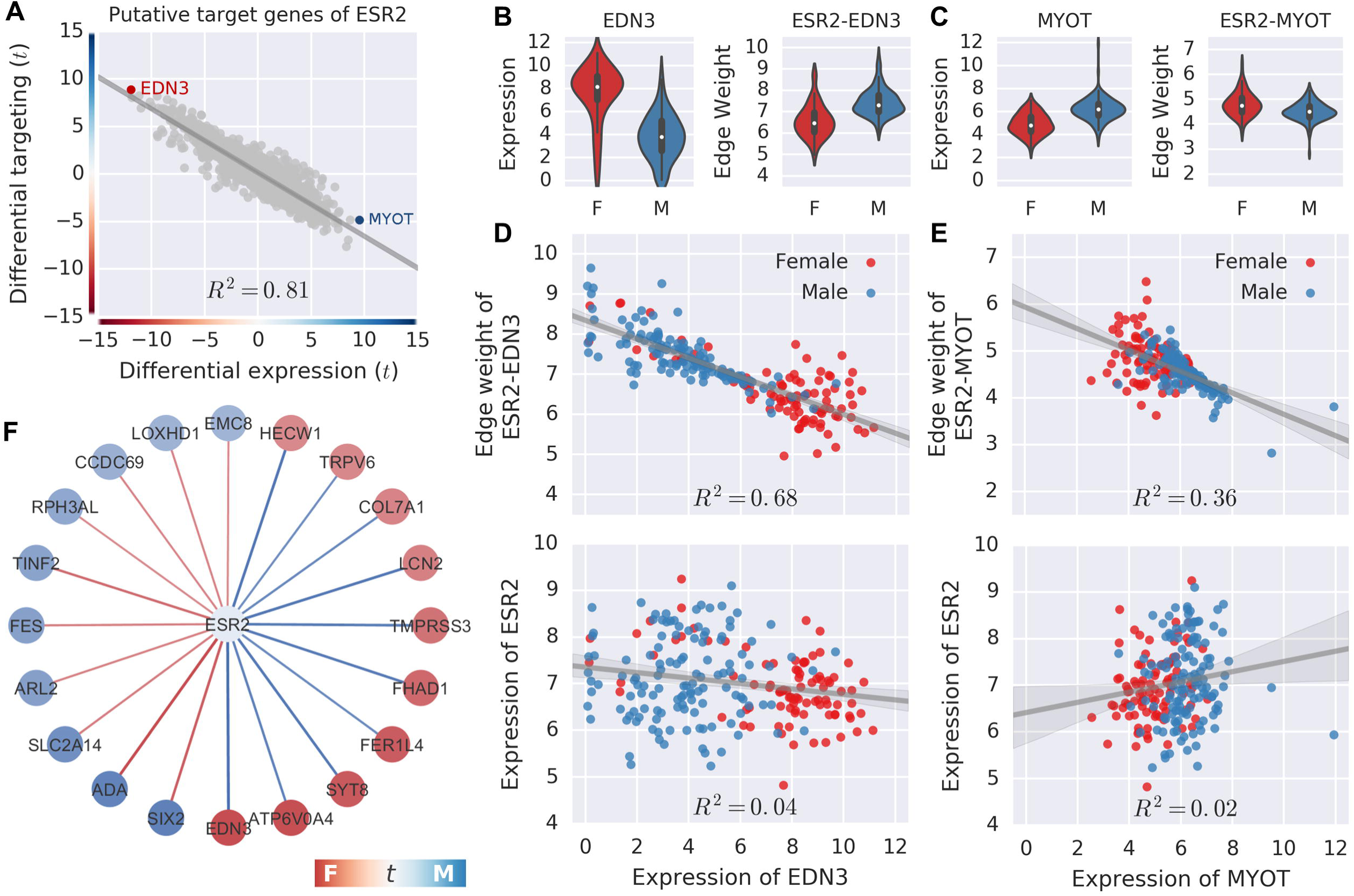
Network inference suggests regulatory potential of transcription factor ESR2 in driving the sexually dimorphic gene expression of its putative target genes, related to Figure 3. **(A)** Inferring sex differential targeting of transcription factor ESR2 on its putative target genes versus differential expression levels. **(B-C)** Comparison of gene expression levels and inferred ESR2 targeting edge weights between females and males for EDN3 **(B)** and MYOT **(C)**, as highlighted in **(A). (D-E)** Expression of EDN3 **(D)** and MYOT **(E)** are highly correlated with ESR2 regulation (upper) but not with ESR2 expression (bottom). **(F)** Network visualization of the top 20 significantly differentially targeted genes of ESR2.

